# Experimental evolution reveals the genetic basis and systems biology of superoxide stress tolerance

**DOI:** 10.1101/749887

**Authors:** Justin Tan, Connor A. Olson, Joon Ho Park, Anand V. Sastry, Patrick V. Phaneuf, Laurence Yang, Richard Szubin, Ying Hefner, Adam M. Feist, Bernhard O. Palsson

**Author notes:** Corresponding author: Bernhard O. Palsson. Author contributions J.T, C.A.O., J.H.P, A.M.F, and B.O.P. designed the study. J.T., J.H.P, R.S., and Y.H. performed experiments. J.T., A.V.S., and P.V.P. analyzed the data. J.T. and B.O.P. wrote the manuscript, with contributions from all the other co-authors.

## Abstract

Bacterial response to oxidative stress is of fundamental importance. Oxidative stresses are endogenous, such as reactive oxidative species (ROS) production during respiration, or exogenous in industrial biotechnology, due to culture conditions or product toxicity. The immune system inflicts strong ROS stress on invading pathogens. In this study we make use of Adaptive Laboratory Evolution (ALE) to generate two independent lineages of *Escherichia coli* with increased tolerance to superoxide stress by up to 500% compared to wild type. We found: 1) that the use of ALE reveals the genetic basis for and systems biology of ROS tolerance, 2) that there are only 6 and 7 mutations, respectively, in each lineage, five of which reproducibly occurred in the same genes (iron-sulfur cluster regulator *iscR*, putative iron-sulfur repair protein *ygfZ*, pyruvate dehydrogenase subunit E *aceE*, succinate dehydrogenase *sucA*, and glutamine tRNA *glnX)*, and 3) that the transcriptome of the strain lineages exhibits two different routes of tolerance: the direct mitigation and repair of ROS damage and the up-regulation of cell motility and swarming genes mediated through phosphate starvation, which has been linked to biofilm formation and aggregation. These two transcriptomic responses can be interpreted as ‘flight’ and ‘fight’ phenotypes.

**Importance:** Bacteria encounter oxidative stress from multiple sources. During pathogenic infections, our body’s immune system releases ROS as a form of antimicrobial defense whilst bacteria used in industrial biotechnology are frequently exposed to genetic modifications and culture conditions which induce oxidative stress. In order to get around the body’s defences, pathogens have developed various adaptations to tolerate high levels of ROS, and these adaptive mechanisms are not always well understood. At the same time, there is a need to improve oxidative stress tolerance for industrially relevant strains in order to increase robustness and productivity. In this study we generate two strains of superoxide tolerant *Escherichia coli* and identify several adaptive mechanisms. These findings can be directly applied to improve production strain fitness in an industrial setting. They also provide insight into potential virulence factors in other pathogens, highlighting potential targets for antimicrobial compounds.

## Introduction

During aerobic respiration, leakage of high energy electrons from respiratory quinones and electron transport chain components creates reactive oxidative species (ROS), such as superoxide and peroxide radicals. Studies have placed the production rate of hydrogen peroxide during normal aerobic growth in *E. coli* to be as high as 10-15μM/s (Imlay 2015; Seaver and Imlay 2004). In addition to the endogenous production of ROS, bacteria also experience oxidative stress from external sources such as the host immune response during pathogenic infections (Kohanski et al. 2007). Production strains of *E. coli* also tend to experience redox stress due to high oxygen concentrations used in fermentation vessels (Baez and Shiloach 2013), or after genetic modifications to the host cell such as the knockout of thioredoxin and glutathione biosynthesis genes (Bessette et al. 1999).

Due to their reactivity, ROS cause damage to macromolecules in the cell. DNA damage is caused by direct oxidation of individual bases and cross-linking between strands (Cadet and Wagner 2013). Lipids are peroxidized, changing membrane permeability. ROS damage to proteins occurs via direct oxidation of vulnerable amino acids such as cysteine and methionine, and loss of metal cofactors (Birben et al. 2012). A major target of ROS damage are iron-sulfur clusters, found in important catalytic proteins and vital for central carbon metabolism and amino acid biosynthesis (Imlay 2006; Roche et al. 2013). Iron-sulfur clusters readily change redox states (Broderick 2003), making them particularly vulnerable to damage by ROS: an initial loss of an electron causes destabilization and a subsequent loss of an iron ion (Djaman, Outten, and Imlay 2004), with further oxidation causing a complete degradation and loss of the iron-sulfur cluster. *E. coli* relies on two systems, the Isc and Suf cluster assembly systems, to synthesize and repair damage to iron-sulfur clusters (Djaman, Outten, and Imlay 2004).

Free cytoplasmic iron is a double-edged sword during oxidative stress. On the one hand, it is used as the metal cofactor in superoxide dismutase *sodB*, which scavenges superoxide species; on the other, it participates in the Fenton reaction, which converts superoxide and peroxide molecules into the even more reactive hydroxyl radical (Farr and Kogoma 1991). Damage to iron-sulfur clusters caused by oxidative stress not only inactivates the iron-sulfur containing protein, but also releases iron into the cytoplasm. Fine control of iron concentrations in the cytoplasm is thus vital to adequately deal with oxidative stress. *E. coli* does this via the transcriptional regulator Fur which represses iron uptake systems when bound to free Fe^2+^. Oxidative stress has been shown to upregulate Fur, resulting in the repression of iron-uptake systems and a reduction of iron levels in wild type cells (Zheng et al. 1999).

Various methods have been explored in order to increase tolerance of *E. coli* to exogenous oxidative stress. Those that were successful made use of mutations to CRP (Basak and Jiang 2012), or the overexpression of damage mitigation genes such as *sodC* and *katG* (Battistoni et al. 2000; Smith, Imlay, and Mackie 2003). *E. coli* also exhibits a phenomenon known as cross-tolerance where various stressors such as cold shock and osmotic shock have also been found to induce the oxidative stress response (Smirnova, Muzyka, and Oktyabrsky 2000; Smirnova, Zakirova, and Oktiabr’skiĭ 2001). When previously exposed to sublethal levels of one type of stress, *E. coli* has shown improved tolerance to a subsequent exposure to lethal levels of another type of stress (Rodríguez-Rojas et al. 2019).

Due to its role in pathogenesis and importance in industrial biotechnology, understanding tolerance to ROS remains a significant area of research. In this paper we make use of tolerization adaptive laboratory evolution (TALE) (Mohamed et al. 2017) to increase the tolerance of *E. coli* to the redox-cycling compound paraquat. We then make use of genome resequencing, transcriptomics, and ribosome profiling to gain insight into these adaptations and the mechanisms through which they increase tolerance.

## Results

### TALE increased tolerance to oxidative stress

Paraquat was chosen as the ROS stressor in these experiments since it is a known generator of internal superoxide stress through redox cycling. In order to differentiate mutations arising from adaptation to the culture media from those arising from oxidative stress, we make use of an *E. coli* K-12 MG1655 strain previously evolved on glucose minimal media as the starting strain (LaCroix et al. 2015). Exponentially growing cultures were passed into incrementally higher paraquat concentrations to increase the tolerance to superoxide stress (Figure 1A).Two replicate end points (PQ1 and PQ2) were generated from the TALE which took place over 30 days, representing over 8 × 10^11^ cumulative cell divisions (CCD) (Lee et al. 2011) (Figure 1B).

**Figure 1.**
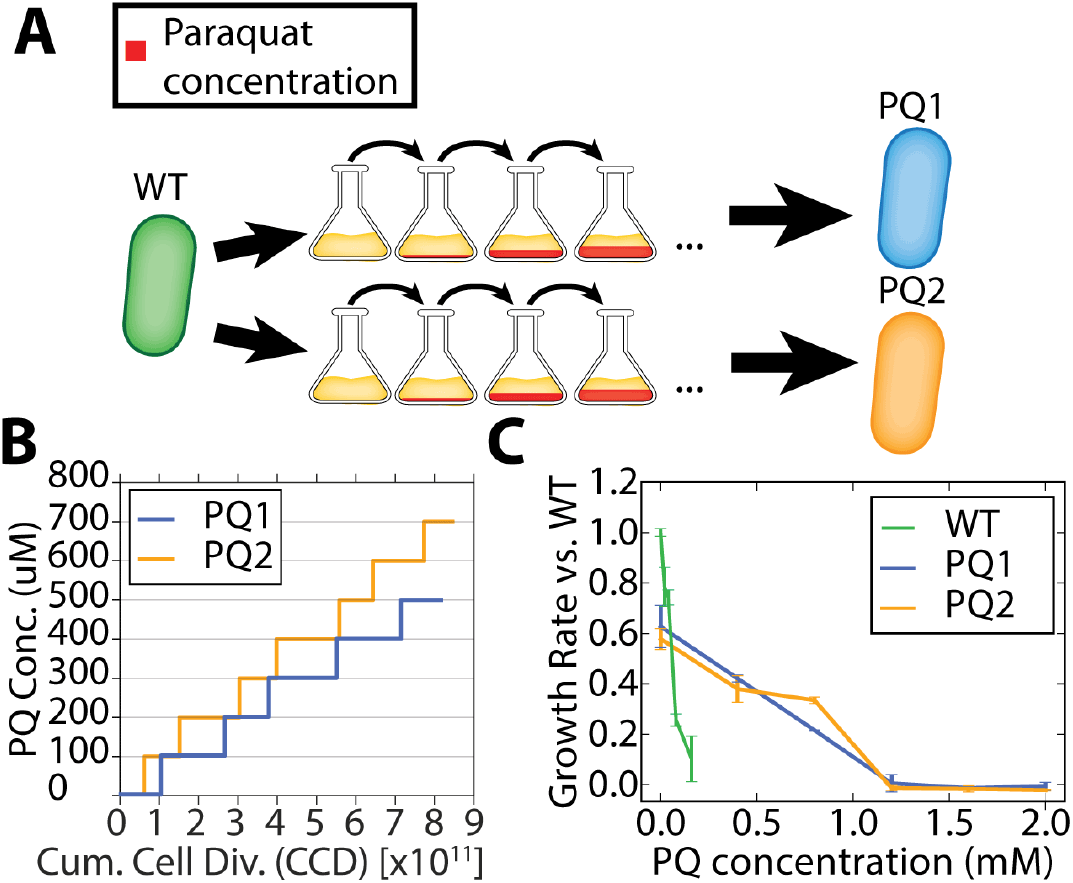
Description of the ALE experiment run and physiological characterization of the evolved strains. A: TALE was used to increase tolerance to paraquat. Paraquat concentration was raised in small increments over the course of evolution, resulting in two end point strains. B: Graph showing paraquat concentration used in TALE against the number of cumulative cell divisions. C: Growth characterization of end point strains. WT shows loss of viability above 0.16mM, while end point strains show loss of viability at 0.8mM.

To quantify the increase to paraquat tolerance, we performed growth experiments of the evolved strains across a range of paraquat concentrations between 0mM and 2.0mM. Wild type showed loss of viability at 0.16mM, while PQ1 and PQ2 only lost viability at paraquat concentrations above 0.8mM, a tolerance increase of 500% (Figure 1C).

### Whole genome resequencing and mutation analysis reveal the genetic basis for increased ROS tolerance

Clonal isolates of the end point populations (PQ1, PQ2) were subject to whole genome resequencing in order to identify the genetic basis of tolerance to paraquat stress. Genetic mutations were called using the *breseq* computational pipeline (Materials and Methods). Key mutations related to tolerance to oxidative stress were determined by identifying genes or genetic regions that were mutated across multiple isolates from independent samples (Table 1). The full list of mutations is deposited in ALEdb (Phaneuf et al. 2019). Only six and seven mutations for PQ1 and PQ2, respectively, were required to confer tolerance to five times the maximum concentration of paraquat compared to WT.

**Table 1.** Mutations found in evolved strains following TALE to increase tolerance to paraquat

| Category | Gene | Strain | Mutation |
| --- | --- | --- | --- |
| Carbon flux | *aceE* | PQ1 | Q791* |
|  |  | PQ2 | Q409* |
|  | *sucA* | PQ1 | R182L |
|  |  | PQ2 | G594S |
|  | *gltA* | PQ2 | L8P |
| Iron-sulfur cluster | *ygfZ* | PQ1 | V107E |
|  |  | PQ2 | T108P |
|  | *iscR* | PQ1 | V55L |
|  |  | PQ2 | C104S |
| tRNA | *glnX* | PQ1 | (C-> T 35/75nt) |
|  |  | PQ2 | (C-> T 35/75nt) |
| Phosphate uptake | *pitA* | PQ1 | INS (+T 5/1500nt) |
| Polymyxin resistance | *arnA* | PQ2 | A21A |

Five genes were mutated in common: *aceE*, *ygfZ*, *iscR*, *sucA*, and *glnX* (Table 1). Two of these five genes (*aceE* and *sucA)* are related to carbon metabolism and the TCA cycle. *aceE* was one of the first genes to be mutated across both replicates, with both independent mutations resulting in truncation of the gene, at residues 791 and 409 for PQ1 and PQ2, respectively. *aceE* encodes the E1 subunit of the pyruvate dehydrogenase complex which catalyzes the reaction that converts pyruvate to acetyl-CoA, a key step for the entry of carbon flux into the TCA cycle during aerobic growth on glucose as the sole carbon source. *sucA*, on the other hand, encodes the E1 subunit of the 2-oxoglutarate dehydrogenase complex that catalyzes the conversion of 2-oxoglutarate into succinyl-CoA and CO_2_ along with the production of NADH. The TCA cycle is the main source of high energy electrons during aerobic respiration, and both these mutations would result in a reduction in carbon flux through the TCA cycle, reducing redox load. Interestingly, both *sucA* and *aceE* are the thiamine-binding E1 components of their respective dehydrogenase complexes, suggesting a significant vulnerability in thiamine production under oxidative stress. Indeed, 2-iminoacetate synthase, coded for by the gene *thiH*, is a key enzyme in the thiamine-biosynthetic pathway and was found to contain a redox sensitive iron-sulfur cluster (Ayala-Castro, Saini, and Outten 2008).

Two other commonly mutated genes are related to iron-sulfur cluster synthesis and repair (*iscR* and *ygfZ*). *iscR* is the transcriptional regulator for iron-sulfur cluster and biosynthesis and is involved in the regulation of the *isc* and *suf* operons (Schwartz et al. 2001). The DNA binding affinity of IscR is dependent on the presence of iron-sulfur clusters bound to the protein. The *icsR* mutation site C104S in PQ2 is known to be one of the three conserved cysteine residues involved in iron-sulfur cluster binding (Fleischhacker et al. 2012), and mutations at this location have been shown to have an effect on the regulation of the *iscRSUA* and *sufABCDSE* operons (Rajagopalan et al. 2013). *ygfZ* was also found to be mutated in both PQ1 and PQ2 at amino acid residue positions 107 and 108, respectively. Though the function of *ygfZ* in *E. coli* is still unclear, it has been shown to be a folate-binding enzyme potentially involved in either the synthesis or repair of iron-sulfur cluster proteins (Jeffrey C. Waller et al. 2010; J. C. Waller et al. 2012). *ygfZ* has been hypothesized to also have a direct role in the degradation of plumbagin (a redox stress causing compound) (Lin et al. 2010), while inactivation of *ygfZ* has been found to result in increased sensitivity to oxidative stress in *E. coli* (Jeffrey C. Waller et al. 2010). Positions 107 and 108 have been previously determined to be conserved across *E. coli*, *M. tuberculosis*, and *K. pneumoniae* (Lin et al. 2010) suggesting that this region might be critical to improving or modifying one or both of the hypothesized functions of *ygfZ* (Fe-S repair and plumbagin degradation).

The last common mutation was found in *glnX*. *glnX* is one of the four glutamine tRNAs in *E. coli* which decodes the CAG codon in wild type. This mutation occurred at nucleotide 35 in the gene, changing the sequence of the anticodon from CTG to CTA, allowing the suppression of the amber stop codon TAG.

The end points also contained one and two strain specific mutations, respectively. PQ1 contains a frameshift mutation in the *pitA* gene, encoding the low affinity phosphate transporter (Harris et al. 2001). *pitA* is the major route for phosphate uptake in the cell under phosphate replete conditions, and its disruption would severely limit phosphate availability in PQ1. PQ2 contains an additional mutation in *gltA* (encoding citrate synthase), another member of the TCA cycle, and a silent mutation in *arnA*, a protein involved in polymyxin resistance. These mutations occur close to the 5’ end of the gene at residue 8 and 21 respectively, which has been shown to affect gene expression levels through changes to mRNA secondary structure (Welch et al. 2009)

### Knockout of aceE improves fitness at low levels of ROS stress

One of the first mutations to show up in both replicates during the first phase of tolerization was a truncation mutation at residue 409 and 791 for PQ1 and PQ2 respectively, suggesting that the disruption of *aceE* activity was important for the resistance to ROS stress. To better understand the role of the aceE truncation in improving strain fitness, we knocked out *aceE* in the wild type strain. The loss of *aceE* had a negative impact on strain viability in M9 minimal media with glucose as the sole carbon source, necessitating the addition of 10% LB to the media which greatly improved tolerance to paraquat, so paraquat tolerance is not directly comparable to the evolved strains. However, we see that compared to WT grown in the same media, Δ*aceE* shows improved fitness when concentration of paraquat is low, but loses this fitness advantage at higher levels of paraquat (Supplementary Figure 1).

### *glnX* mutation affects *aceE* expression

The truncation mutation that occurred in both replicates makes use of a TAG stop codon, and both strains then independently developed a mutation in *glnX* that allows suppression of the TAG stop codon. Stop codon suppression is known to be incomplete owing to competition between the native ribosome release factor and the suppression tRNA (Eggertsson and Söll 1988). In order to see the effect of the *glnX* suppressor mutation on the expression level of *aceE*, we make use of ribosome profiling to determine the ribosomal density on the *aceE* gene. Due to the great selective disadvantage of the loss of *aceE*, we were unable to revive the PQ1 midpoint, and hence are looking only at the PQ2 midpoint. We calculated the ratio of the ribosomal density before and after the truncation to determine the percentage of stop codon readthrough. In WT, ribosome density downstream of the truncation location (Q409*) is almost equal to that found upstream (0.97±0.11). With the non-sense mutation but without the *glnX* suppressor mutation, this ratio drops to 0.05±0.01 due to premature truncation of translation at the non-sense mutation. In PQ2 with the *glnX* suppressor mutation, we see that the ratio increases to 0.23±0.08, indicating that the *glnX* mutation serves to tune the translation level of *aceE* (Figure 2B).

**Figure 2.**
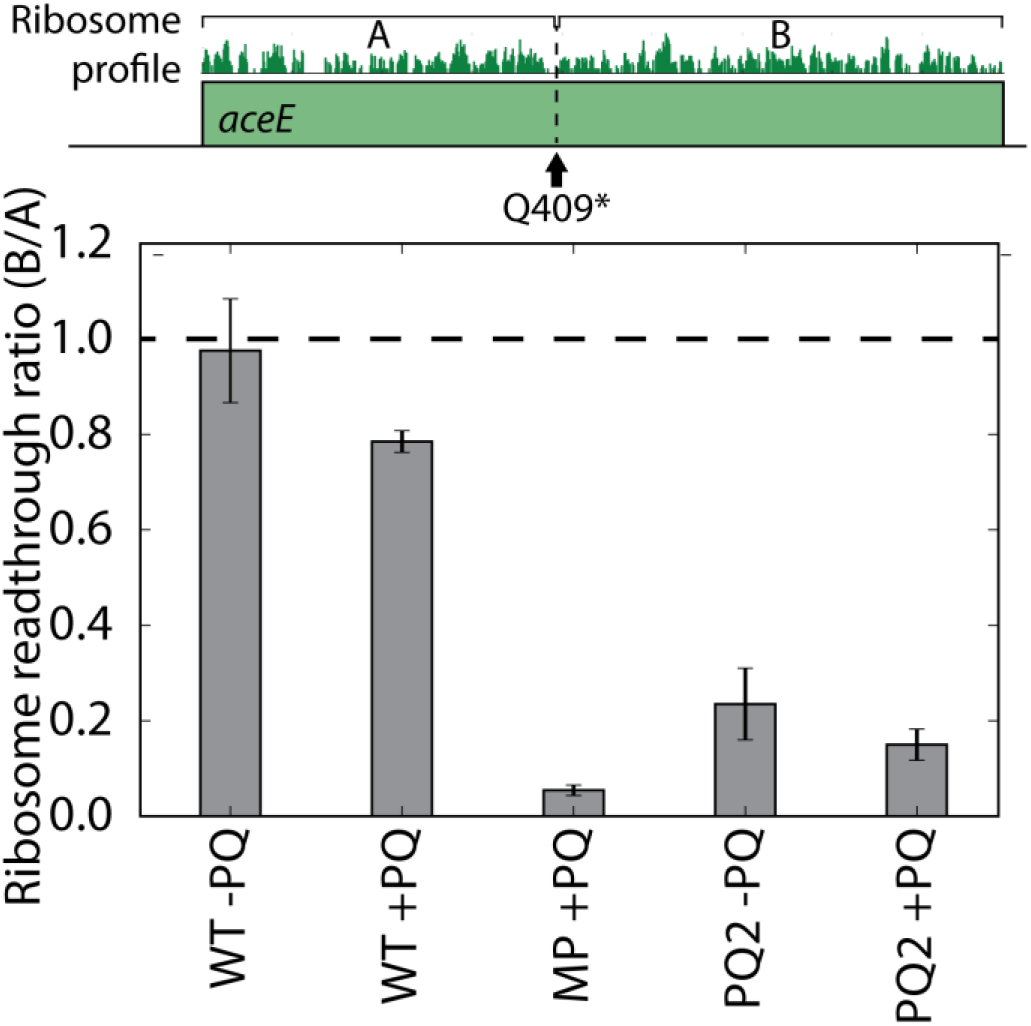
The *glnX* suppressor mutation allows limited readthrough of the non-sense mutation. We calculate the ratio of ribosomal density downstream and upstream of the non-sense mutation. WT shows readthrough ratios close to 1.0, indicating that there is no truncation along the *aceE* gene. Conversely, a midpoint clonal isolate with the *aceE* non-sense mutation but without the *glnX* suppressor mutation shows a very low ratio, suggesting that translation terminates at the non-sense mutation. PQ2 end point strains with both the *aceE* non-sense mutation and the *glnX* suppressor mutation show increased readthrough of the non-sense mutation, suggesting the combination of these mutations result in a tuning of aceE levels in the cell.

### Dysregulation of iron-uptake genes under stress

We calculated differential gene expression of the evolved strains PQ1 and PQ2 when subject to 0.25mM of paraquat for 20 minutes relative to no paraquat exposure. The response of WT *E. coli* to superoxide stress has previously been well-characterized, involving the up-regulation of the SoxRS and OxyR regulons (Seo et al. 2015; Farr and Kogoma 1991; Rui et al. 2010), thus we subtracted the ROS stress response in WT from the set of differentially expressed genes in the evolved strains in order to isolate adaptation specific transcriptional responses. This process left us with a total of 49 DEGs in PQ1 (47 up-regulated, and 2 down-regulated) and 75 DEGs in PQ2 (68 up-regulated, and 7 down-regulated), that were not part of the canonical superoxide stress response in WT. We find that both PQ1 and PQ2 have a convergent transcriptional response to ROS stress. Also in both, 45 genes were up-regulated and 2 genes were down-regulated. The most differentially regulated COG categories during stress were secondary metabolites (PQ1 p-value = 3.17×10^−12^, PQ2 p-value = 8.64×10^−10^) and Inorganic ion transport (PQ1 p-value = 2.04×10^−16^, PQ2 p-value = 1.71×10^−18^) (Supplementary Figure 2EFGH, Table 2).

**Table 2.** Important differentially regulated genes with and without paraquat stress. In the unstressed condition, differential expression of evolved strains was calculated relative to WT strain. In stressed conditions, differential expression of evolved strains under paraquat stress was calculated relative to the same strain without paraquat stress. Genes differentially regulated in WT under paraquat stress were subtracted to isolate evolved strain specific adaptations.

| Condition | Strain | Function | Genes | Regulation |
| --- | --- | --- | --- | --- |
| Unstressed | Common | Not well characterized/<br>non-functional | *ygfZ, ybckLM, bcsAB* | Up-regulated |
|  |  | Osmotic stress | *proVWX* | Up-regulated |
|  |  | Fatty acid degradation | *fadAB* | Up-regulated |
|  |  | Sulfur uptake | *tauABC* | Down-regulated |
|  | PQ1 | Phosphate homeostasis | *pstABC, phoU, phoBR* | Up-regulated |
|  |  | Cell motility | *fliHIJKMOPR, flhABE, motAB, flgABEFGHIKLMN, fliACDEFGNSTZ, trg, cheAZYBRW, dgcZ, tar, tsr, tap* | Up-regulated |
|  | PQ2 | Iron-sulfur cluster repair and synthesis | *iscASRU, sufABCDES, fdx, hscAB* | Up-regulated |
|  |  | DNA synthesis | *nrdHIEF* | Up-regulated |
|  |  | Cell motility | *fimE* | Up-regulated |
|  |  |  | *cheAWR, motAB, tap, tar, fimABCDFGI* | Down-regulated |
|  |  | Fermentative respiration | *ldhA, pta, ackA* | Up-regulated |
|  |  | TCA cycle | *sdhBCD* | Down-regulated |
| Stressed | Common | Iron uptake and storage | *entSABCDEFH, fepABCDGRI, yncD, fes, fiu, fhuE* | Up-regulated |
|  |  | DNA synthesis | *nrdEFH* | Up-regulated |
|  |  | DNA repair | *ligA* | Up-regulated |
|  |  | Iron-sulfur cluster | *sufDES* |  |
|  | PQ1 | Iron-sulfur cluster repair and synthesis | *iscR* | Up-regulated |
|  | PQ2 | Formate dehydrogenase | *fdnH, fdoG* | Up-regulated |
|  |  | Motility | *ybjN* | Up-regulated |
|  |  | marR operon | *marA, mdaB* | Up-regulated |
|  |  | ROS scavenging | *sodB* | Down-regulated |

These 45 genes up-regulated in common are highly enriched for regulation by Fur (p-value = 8.91 × 10^−27^), and almost all of them are directly related to the uptake and storage of iron. In contrast, we see that WT actually down-regulates these genes, despite all three strains directly upregulating Fur (Figure 3). During superoxide stress, free iron in the cell aggravates damage to cellular components through formation of hydroxyl radicals via the Fenton reaction (Winterbourn 1995). As the regulatory activity of Fur is mediated by availability of Fe^2+^, this could indicate differences in the availability of free iron in the cell. Other commonly upregulated genes during stress are related to DNA synthesis and repair, such as ribonucleotide reductase *nrdEFH* and DNA ligase *ligA*, which would allow cells to repair DNA damaged by ROS at an increased rate compared to WT.

**Figure 3.**
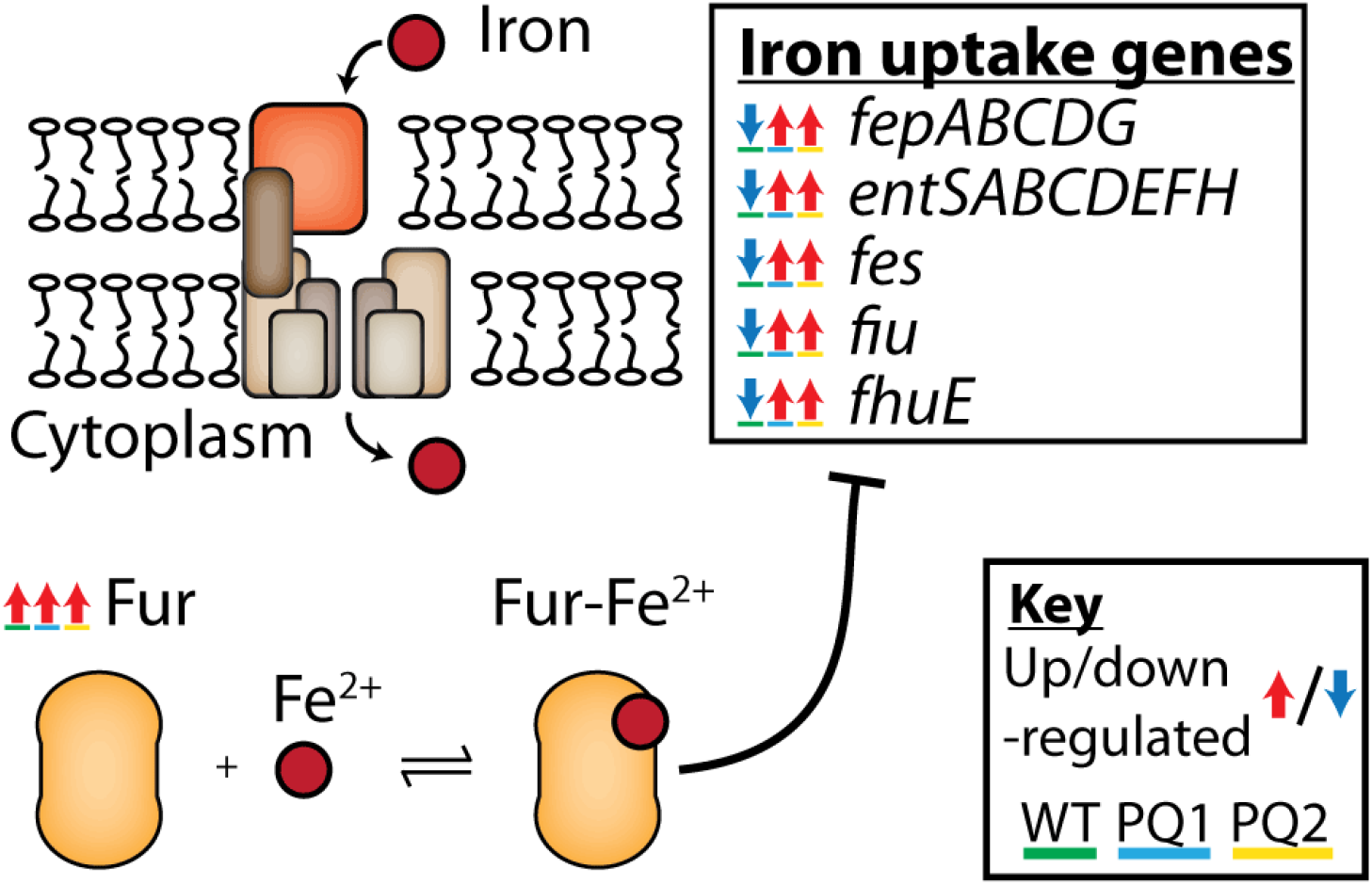
Evolved strains show dysregulation of iron-uptake genes regulated by Fur-Fe^2+^ while under paraquat stress. Differential gene expression was calculated for each strain under paraquat stress relative to the same strain under normal growth conditions; significantly differentially expressed genes are circled in bold. Both WT and the evolved strains up-regulate Fur, which in WT proceeds to repress the iron-uptake genes below. In contrast, the evolved strains instead see an up-regulation of these iron-uptake genes.

### Transcriptomic characterization of end point strains under normal growth conditions reveals two ROS tolerant phenotypes

When grown without paraquat stress, we find that PQ1 and PQ2 differentially express 126 and 182 genes, respectively, relative to wild type (PQ1: 104 up-regulated, 22 down-regulated, PQ2: 89 up-regulated, 92 down-regulated). Of these, only 15 genes are commonly up-regulated and 11 genes are commonly down-regulated (Table 2, Supplementary Figure 2AB). PQ1 significantly up-regulates while PQ2 significantly down-regulates genes with the COG annotation of Cell motility (PQ1 p-value = 4.71×10^−94^, PQ2 p-value = 1.43×10^−17^), while PQ2 also differentially regulates genes in the COG category of Energy production (p-values = 1.02×10^−5^) (Supplementary Figure 2CD).

Several of the commonly upregulated genes, *ygfZ*, *ybcKLM*, and *bcsAB*, are not well characterized or thought to be non-functional in K-12 strains. *ygfZ*, discussed above, is annotated as a putative iron-sulfur cluster repair protein and is both mutated and up-regulated in PQ1 and PQ2 (Jeffrey C. Waller et al. 2010). The combination of the mutation and up-regulation of *ygfZ* indicates that it plays an important role in tolerance to paraquat stress and warrants further investigation. *ybcL*, which is in an operon with *ybcM* directly downstream of *ybcK*, has previously been found to inhibit neutrophil migration during bladder infections by the pathogenic strain of *E. coli* UTI89, although several mutations in the K-12 variant have caused a loss of its original pathogenic function (Lau, Loughman, and Hunstad 2012). Regulation of the *ybcLM* operon is not well understood, and up-regulation of these genes in the evolved strains suggests a vestigial virulent response. *bcsA* and *bcsB* synthesize cellulose in non-K12 *E. coli* strains, but in K-12 strains a nonsense SNP in upstream gene *bcsQ* results in decreased expression of *bcsA* (Serra, Richter, and Hengge 2013).

The evolved strains also commonly up-regulate the genes *proVWX* which are related to tolerance to osmotic stress. Previous studies have shown that previous exposure to one type of stress, such as pH, osmotic, oxidative, or thermal stress, has a positive effect on culture tolerance to other stresses, possibly due to the induction of general stress response pathways through RpoS (Rodríguez-Rojas et al. 2019). Evolution of these strains in the presence of oxidative stress might cause a constitutive upregulation of stress response pathways, resulting in upregulation of *proVWX* even in the absence of paraquat.

Two genes utilized in fatty acid degradation, *fadAB*, are commonly upregulated. The truncation of *aceE* would result in a decrease of conversion from pyruvate to acetyl-CoA, lowering the available pool of acetyl-CoA which is important for synthesis of various amino acids through the glyoxylate shunt and TCA cycle intermediates. Upregulation of the fatty acid degradation pathway would increase levels of acetyl-CoA in the cell through oxidation of fatty acids.

### PQ1 – Exhibits a “Flight” Phenotype

As mentioned above, the frame shift mutation in *pitA* disrupts its function, severely limiting phosphate uptake in PQ1 (Rosenberg, Gerdes, and Chegwidden 1977). Expectedly, PQ1 up-regulates the high affinity phosphate transport complex genes *pstABCS* and phosphate regulators *phoBR* and *phoU* even when not under paraquat stress, suggesting that the cell is constitutively experiencing phosphate starvation. Phosphate limitation is observed to activate a swarming phenotype in *P. aeruginosa* (Bains, Fernández, and Hancock 2012). We observe the similar up-regulation of cell motility related genes in PQ1. 42 of the 126 differentially regulated genes in PQ1 are related to cell motility. These genes include flagellar biosynthesis genes *fliOIHKJPRM and flhABE*, flagellar components *motAB*, *flgABEFGHIKLMN*, and *fliACDEFGNSTZ*, chemotaxis related proteins *trg* and *cheAZYBRW*, as well as motility regulators *dgcZ, tar, tsr*, and *tap*. The up-regulation of these genes has also been associated with auto-aggregation and biofilm formation, which has been shown to increase the tolerance of *E. coli* to various stresses including oxidative stress (Laganenka, Colin, and Sourjik 2016).

Surprisingly, this regulatory adaptation in PQ1 does not intuitively counter ROS stress. Instead, it increased cellular motility (Supplementary Figure 3), which might lead to a secondary effect of oxidative stress tolerance. We therefore term this the “Flight” phenotype (Figure 4)

**Figure 4.**
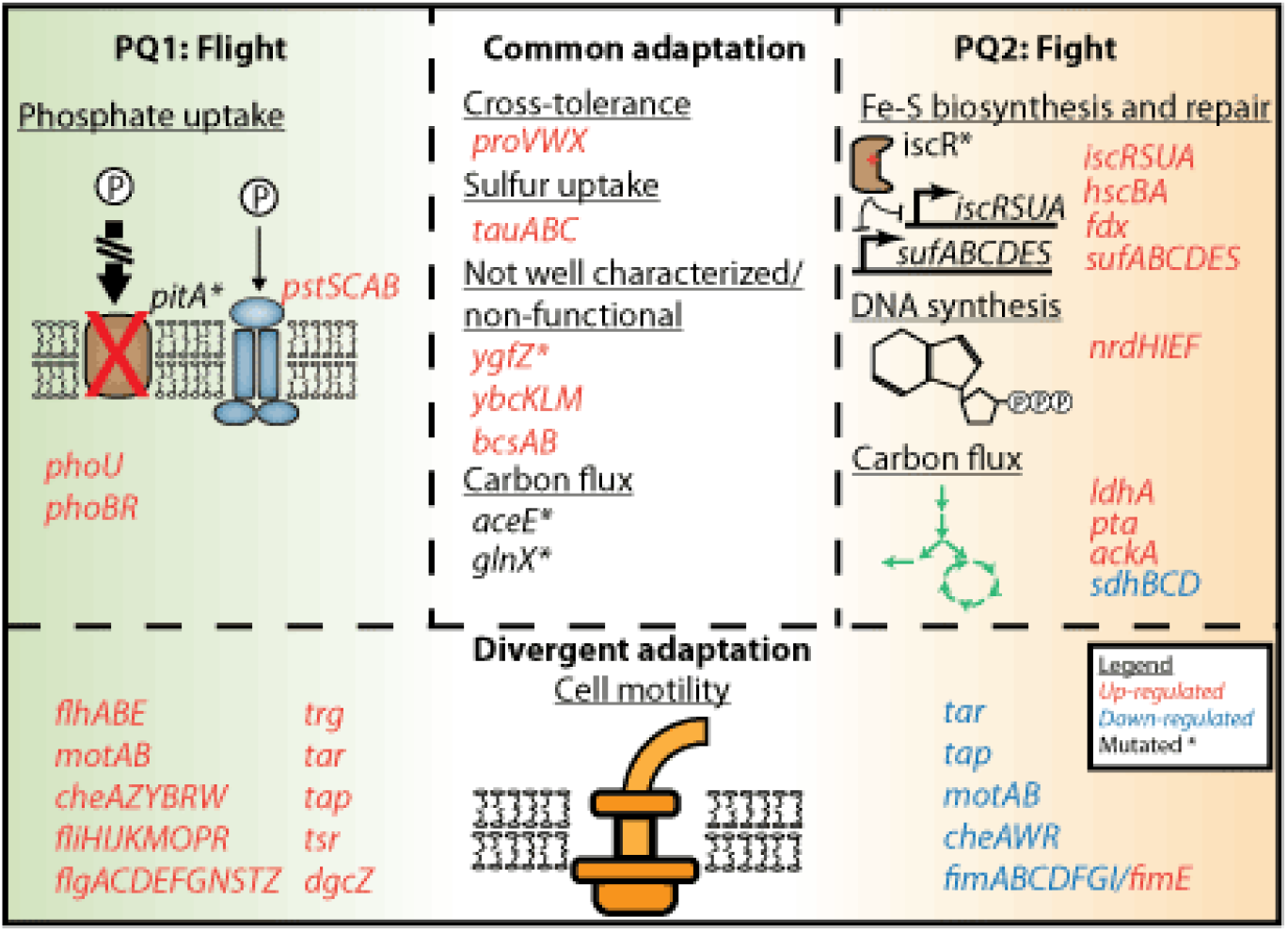
PQ1 and PQ2 show two different phenotypes growing under normal conditions. Differential expression is calculated for the evolved strains growing without paraquat stress relative to wild type. Up-regulated genes are highlighted in red, down-regulated genes are highlighted in blue, mutated genes are denoted by an asterisk. The “Flight” phenotype used by PQ1 makes use of a *pitA* disruption to induce phosphate starvation, providing a regulatory impetus for increased cell motility and aggregation, increasing tolerance to oxidative stress. On the other hand, PQ2 exhibits the “Fight” phenotype wherein it up-regulates genes involved in the repair moieties damaged by ROS, such as iron-sulfur clusters and DNA, as well as redirects carbon flux from the TCA cycle towards fermentative pathways.

### PQ2 – Exhibits a “Fight” Phenotype

In contrast to PQ1, PQ2 downregulates cell motility related genes *cheAWR*, *motAB*, *tap*, and *tar*, and fimbriae genes *fimABCDFGI*, while up-regulating *fimE*. These changes to the FimB/FimE ratio are consistent with the overall downregulation of type 1 fimbriae expression (Holden et al. 2007; Klemm 1986).

We find that the mutation in the iron-sulfur cluster binding region of the protein IscR causes a dysregulation of its regulon. Genes regulated by iscR, such as *iscASRU*, *hscSB*, *fdx*, *sufABCDES*, and *nrdHIEF* are all up-regulated in PQ2 even in the absence of paraquat stress. These genes synthesize and repair iron-sulfur clusters and DNA, which might increase the rate of cellular repair of ROS damage. Interestingly, *nrdEF* has been found to only be induced under iron starvation (J. E. Martin and Imlay 2011; Andrews 2011), providing further support that iron levels in the evolved strains might differ from wild type, leading to the differences found in regulation of iron uptake genes.

In addition to mutations found in *aceE*, *gltA*, and *sucA* genes, PQ2 also upregulates *ldhA*, *pta*, and *ackA*, indicating a switch from aerobic respiration to fermentation for the production of lactate or acetate. Succinate dehydrogenase subunits *sdhBCD* are also downregulated, further decreasing flux through the oxidative arm of the TCA cycle.

Overall, PQ2 seems to adopt a different tolerance strategy from PQ1: it up-regulates genes that allow it to repair the damage caused by ROS to iron-sulfur clusters and DNA, and changes the flow of carbon flux towards fermentative pathways to mitigate endogenous and paraquat cycling ROS production. We have accordingly termed it the “Fight” phenotype (Figure 4).

## Discussion

Bacteria encounter and respond to oxidative stress in many environments, both while fighting an immune response and responding to culture conditions in industrial biotechnology (Baez and Shiloach 2013; Bessette et al. 1999). Our results reveal a relatively simple genetic basis for adaptation of *E. coli* to extremely high levels of superoxide stress. They also uncover several novel genetic adaptations and regulatory strategies used by *E. coli* for adaptation to oxidative stress, including the use of a non-sense/suppressor mutation pair to control translation of pyruvate dehydrogenase in order to reduce carbon flux into the TCA cycle; dysregulation of the Fur regulon; phosphate starvation as a regulatory impetus for oxidative stress tolerance; and direct mitigation of ROS damage through. These findings pave the way to a better understanding of host-pathogen interactions as well as suggesting avenues for designing host strains to better withstand oxidative stress.

Over the course of this study, we have generated several significant findings. First, the use of experimental evolution was successful in developing strains that were tolerant to increased levels of paraquat. *E. coli* can develop up to a 500% increase in tolerance to paraquat relative to wild type in the span of 30 days, with an average of only 6.5 mutations, demonstrating the ease with which *E. coli* adapts to elevated oxidative stress.

Second, the low number of mutations required for adaptation makes the genetic basis of adaptation to oxidative stress simple and interpretable. These mutations fall into two main categories: modulation of metabolic flux through the TCA cycle through the *aceE*, *sucA*, and *glnX* mutations, and the increased synthesis and repair of iron-sulfur clusters through mutations in *ygfZ* and *iscR*. Modulating flux through the TCA cycle is a preventative measure, limiting the availability of high energy electrons which reduces endogenous ROS production and redox cycling by paraquat. On the other hand, mutations that increase synthesis and repair rates of iron-sulfur clusters work to mitigate damage to iron-sulfur clusters caused by ROS. One of the more surprising findings from this study was the occurrence of an amber suppressor mutation in *glnX*, which allowed control of pyruvate dehydrogenase levels in the cell through inefficient readthrough of a non-sense mutation, an evolutionary strategy that has not been previously seen in other experiments. While their role is not well understood, suppressor mutations are commonly found in bacteria that interact directly with hosts, such as those in the gut microbiome (Marshall and Levy 1980), as well as in laboratory strains of *E. coli* that have been subject to long periods of mutagenesis (Belin 2003). These occurrences, as well as the emergence of the suppressor mutation while under oxidative stress, point to the role of suppressor mutations as stress-tolerance mechanisms.

Third, transcriptomics provides useful insight into the tolerization mechanisms of the evolved strains. One interesting finding in this study is the dysregulation of iron-uptake genes. It has long been known that iron plays an important role in oxidative stress in bacteria (Touati 2000). In the presence of oxidative stress, damage to iron-sulfur clusters as a result of the production of ROS results in the release of free iron into the cytoplasm (Djaman, Outten, and Imlay 2004). In WT, Fur binds to free Fe^2+^ repressing the expression of iron-uptake genes in order to control levels of cytoplasmic iron. In the evolved strains, up-regulation of iron-uptake genes instead, in spite of the up-regulation of Fur, might indicate a decreased Fe^2+^ level. This suggests that the evolved strains reduce Fe^2+^ concentration under oxidative stress through a yet to be uncovered mechanism.

Finally, analysis of the evolved strains from a systems biology perspective leads to the identification of two phenotypic states. The first of these makes use of cross-tolerization to phosphate starvation as a means to gain resistance to oxidative stress. The mutation in the *pitA* gene in PQ1 results in a frameshift mutation, disrupting activity of the major phosphate transporter in *E. coli*. The ensuing phosphate starvation phenotype could augment tolerance to oxidative stress in three ways. The first is via the reduction of the availability of high energy electrons for redox cycling and endogenous ROS production (Marzan and Shimizu 2011; Moreau 2004). The second is as a regulatory impetus for up-regulation of general stress response through the RpoS regulon (Mandel and Silhavy 2005). The appearance of the amber suppressor mutation that has previously been observed to relax the stringent response (Breeden and Yarus 1982) might further optimize the response towards oxidative stress. Third, there is evidence that phosphate starvation causes a virulence phenotype which might impart tolerance to oxidative stress as a secondary effect. The *pstABC* complex, upregulated during phosphate starvation, has been found to play an important role in virulence and tolerance to oxidative stress in Avian Pathogenic *E. coli (Crépin et al. 2008)*, while phosphate starvation has been linked to the induction of cellular motility, auto-aggregation, and the formation of biofilms (Vogeleer et al., n.d.), a physical method of increasing resistance to oxidative stress (Schembri et al. 2003). The exposure to low levels of one type of stress resulting in a gain in tolerance to various other seemingly unrelated stressors has previously been observed in several experiments (Gunasekera, Csonka, and Paliy 2008), but this study reveals a potential application of this phenomenon. This result could find a use in metabolic engineering applications where oxidative stress to the cell factory is a concern through simple modifications of phosphate availability in the media.

The second of these phenotypic states increases tolerance to oxidative stress by directly combating the damaging effects of ROS. The mutation in *iscR* in PQ2 causes a constitutive up-regulation of genes in its regulon, such as *iscSUA*, *sufABC*, and *nrdEFH*. The high level of IscSUA and SufABC could confer tolerance to oxidative stress by both increasing the iron-sulfur synthesis rates as well as increasing the rate of repair for damaged iron-sulfur clusters, while NrdEFH might increase DNA synthesis rates in order to mitigate damage done to DNA. In conjunction, a shift from aerobic respiration towards a fermentative mode through coordinated mutations and regulation to reduce metabolic flux through the TCA cycle and up-regulation of genes such as *ldhA*, *ackA*, and *pta* reduces the availability of high energy electrons, reducing the overall production of both endogenous and redox-cycling ROS.

In this study we found that laboratory evolution of *E. coli* leads to adapted strains that can withstand up to five times increased paraquat concentrations compared to wild type. Analysis of these strains revealed insights into the genomic basis for and systems biology of ROS tolerance. Laboratory evolution resulted in adapted strains whose properties were consistent with known targets of ROS damage, yet achieved tolerance through non-intuitive mechanisms. Taken together, the properties of the adapted strains encourage continued work to build a more complete understanding of adaptation to ROS stress.

## Acknowledgements

This work was funded by the Novo Nordisk Foundation Grant Number NNF10CC1016517 and by National Institute of General Medical Science grant number GM057089. We would like to thank Marc Abrams for his assistance with manuscript editing and Amitesh Anand for helpful discussions and assistance with growth screens.

## Methods

### Strains

The initial strain used for the first phase of evolution was an MG1655 K-12 *E. coli* strain which had been evolved for optimal growth on glucose as a carbon source in M9 minimal media (LaCroix et al. 2015).

### TALE

TALE was performed using a similar protocol to that in Mohamad *et. al.* 2017 *(Mohamed et al. 2017)*. Parallel cultures were started in M9 minimal medium by inoculation from isolated colonies. Evolution was performed in an automated platform with 15 mL working volume aerobic cultures maintained at 37°C and magnetically stirred at 1100 rpm. Growth was monitored by periodic measurement of the 600 nanometer optical density (OD600) on a Tecan Sunrise microplate reader, and cultures were passaged to fresh medium during exponential cell growth at an OD600 of approximately 0.3. Growth rates were determined for each batch of medium by linear regression of ln(OD600) versus time. At the time of passage, PQ concentration in the fresh medium batch was automatically increased if a specified growth rate had been met for several flasks. Samples were saved throughout the experiment by mixing equal parts culture and 50% v/v glycerol and storing at −80--°C glycerol.

### Mutation calling

The *breseq* pipeline version 0.33.1 (Deatherage and Barrick 2014) was used to map the DNA-seq reads to an *E. coli* K12 MG1655 reference genome (NCBI accession NC_000913 version 3). DNA-seq quality control was accomplished using the software *AfterQC* version 0.9.7 (Chen et al. 2017).

### Generation of *aceE* knockout

We used P1 phage transduction to transfer the *aceE* knockout from the Δ*aceE* strain in the Keio collection to WT (Baba et al. 2006; Thomason, Costantino, and Court 2007). Briefly, Δ*aceE* strain from the Keio collection was grown up and lysed using P1 phages. The lysate was filtered to remove cell debris leaving behind only P1 phages with packaged Δ*aceE* strain DNA. This was used to infect WT to transfect the DNA. Because the Δ*aceE* strain in the Keio collection contains a selective kanamycin marker at the gene lesion we were able to select for successful transfections.

### Growth curves

End point strains were inoculated from overnight cultures into M9 minimal media with glucose as a carbon source (0.4% w/v) and allowed to grow to A600 ~OD 0.5. They were then diluted down to OD 0.01 with glucose minimal media containing different concentrations of paraquat. These were loaded onto a Bioscreen C set to measure OD every 30 minutes for 24 hours at 37C at high shaking. Comparison between WT and *aceE* knockout was performed in a similar fashion except with 10% LB added to the media to allow growth of the *aceE* knockout strain.

### Culture conditions

WT, PQ1 and PQ2 were grown overnight in M9 minimal media with 0.4% w/v glucose as a carbon source. Fresh media was inoculated with the overnight culture to an initial OD of 0.025. Cultures were aerated with a stir bar in a water bath maintained at 37C until OD reached 0.5. 50mM paraquat was added to a final concentration of 250uM in stressed condition flasks. After 20 minutes both stressed and unstressed conditions were harvested for ribosome profiling and transcriptomics.

### Ribosome profiling

Ribosome profiling libraries were created using a modified version of the protocol outlined in Latif *et. al. (Latif et al. 2014)*. Differences from the published protocol are outlined below. In order to negate the possible confounding effects of addition of chloramphenicol to the media at harvest, cells were lysed by grinding in liquid nitrogen. 50mL of cells were harvested by centrifugation for 4 minutes at 37C in a 50mL conical tube containing 0.400g of sand, supernatant was aspirated quickly and the cell pellet was flash frozen in liquid nitrogen. Pellets were then transferred into a liquid nitrogen cooled mortar and pestle, 500uL of lysis buffer was added and the pellet was pulverised to lyse the cells. Lysate was transferred to a falcon tube to thaw on ice and centrifuged and the supernatant whole cell lysate was isolated to continue with the published protocol (Latif et al. 2014). Reads were sequenced on an Illumina HighSeq machine using a single end 50bp kit.

Ribosome profiling reads had adapters removed using CutAdapt v1.8 (M. Martin 2011), then mapped to *E. coli* genome MG1655 using Bowtie v1.0.0 (Langmead 2010) and scored at the 3’ end to generate ribosome density profiles for each gene.

### Transcriptomics

Cells were pelleted and lysed with a modified version of the RNAProtect Bacteria Reagent protocol, ribosomal RNA was depleted using Ribo-Zero rRNA Removal Kit for Gram-Negative bacteria (Illumina), libraries were created using KAPA RNA Library Preparation kit. Deviations from the kit protocols are mentioned below. 3mL of induced culture was added to 6mL of RNAProtect Bacteria Reagent (Qiagen) and vortexed, then left at room temperature to incubate for 5 minutes. Cells were pelleted and then resuspended in 400uL elution buffer and then split into two tubes, with one kept as a spare. One pellet was then lysed enzymatically with the addition of lysozyme, proteinase-K and 20% SDS. SUPERase-In was added to maintain the integrity of the RNA. RNA isolation was then performed according to the rest of the kit protocol. rRNA was the depleted using the Ribo-Zero rRNA Removal Kit for Gram-Negative Bacteria according to the protocol, and libraries were constructed for paired-end sequencing using the KAPA RNA-Seq Library Preparation kit protocol. Reads were sequenced on the Illumina NextSeq platform.

Transcriptomic reads were mapped using Bowtie2 (Langmead and Salzberg 2012), and reads were counted using HTSeq (Anders, Pyl, and Huber 2010). Differential expression of genes was called using the DESeq2 (Love, Huber, and Anders 2014) package in Bioconductor. Genes with a log_2_fold change greater than 1 and an FDR-adjusted p-value smaller than 0.1 were considered to be significantly differentially expressed between conditions. Raw read counts were normalized to transcripts per million (TPM) for further analysis. Sequencing data is available in the Gene Expression Omnibus (GEO) database with accession number GSE134256.

### Enrichment analysis for COG categories

Differentially expressed genes between pairs of conditions were annotated with their Clusters of Orthologous Genes (COG) categories. We then performed a hypergeometric test to test for enrichment of each COG category amongst the set of differentially regulated genes. The Bonferroni correction was used to adjust for the FDR, and an adjusted p-value below 0.01 was considered significantly enriched.

### Cell motility assay

Overnight cultures of each strain were inoculated into M9 minimal media plates containing 0.4% w/v glucose and 0.25% w/v agar by inserting a pipette tip containing 1uL of culture about 3mm into the center of the plate and ejected as the tip was lifted up. Plates were incubated at 37C for 72 hours.

### Data availability

Genome resequencing data is available at ALEdb.org

RNASeq and Ribosome profiling data is available at the Gene Expression Omnibus (GEO) database using accession number GSE134256

